# Animal-free peptones do not alter bacteriophages propagated for therapeutic use

**DOI:** 10.64898/2026.02.25.707854

**Authors:** Daniel R. Laucirica, Phoebe G. Carr, Mitchell G. Hedges, Andrew Vaitekenas, Zlatibor Velickovic, Stephen M. Stick, Samuel T. Montgomery, Anthony Kicic, Phage WA

## Abstract

**Aims:** Bacteriophage (phage) propagation has traditionally relied on bacterial culture media containing animal-derived ingredients; however, safety concerns with animal-derived materials for production of phages for therapeutic use limit their acceptability. We compared animal-free and traditional media formulations, and evaluated their effects on phage yield, bactericidal activity, and genomic characteristics, hypothesizing no significant differences would be observed.

**Methods and Results:** Phages targeting *Pseudomonas aeruginosa* (n=8) and *Staphylococcus aureus* (n=1) were propagated in solid and liquid media containing animal-free (AF) or animal-derived (LB) peptones. Kinetic assays were used to assess phage suppression of host bacterial growth. In a mock therapeutic phage screen, spot tests, Efficiency of Plating (EOP) and kinetic assays were performed against novel bacterial targets. Whole genome sequencing of phages and their bacterial hosts propagated in AF or LB broth was used to observe genomic differences between formulations. Animal-free peptone did not impact phage yield, with both AF and LB phage stocks growing to high titers (≥10^8^ PFU/mL). Kinetic assay results showed similar suppression indices for AF and LB-grown phages. Likewise, phage screen spot test, EOP, and kinetic assay results were similar between AF and LB phages. Comparisons of phage and bacterial genome annotations showed no major differences arising from media formulation.

**Conclusions:** Findings suggest animal-free peptones do not significantly alter phage yield, bactericidal activity, or genomic characteristics, supporting use of animal-free medium for medicinal phage manufacture. This is one of the first studies to systematically combine phenotypic and genomic assessment of phages and hosts across animal-free and traditional media.

**Impact Statement:** Phage therapy is increasingly used to treat antimicrobial resistance infections. Emerging guidelines and regulations for the manufacture of phage therapeutics will impact laboratory processes and materials used for phage production. Here, we explored the use of an animal-free medium for medicinal phage propagation, providing data on phage yield and metrics of phage activity.

## Introduction

Antimicrobial resistance (AMR) remains a global health threat, with 1.91 million AMR-attributable deaths and 8.22 million AMR-associated deaths predicted to occur by 2050 (G. B. D. Antimicrobial Resistance Collaborators 2024). Bacteriophages or phages, viruses that exclusively infect bacteria, are increasingly used as an experimental alternative therapy for treating antimicrobial resistant infections (Kakasis and Panitsa 2019; Pirnay et al. 2024b). Full translation and clinical implementation of phage therapy remain hindered in part by a lack of quality standards and guidelines for phage manufacture (Ng et al. 2020). While magistral phage preparations have helped formalize frameworks for the creation of individually tailored phage therapeutics (Pirnay and Verbeken 2023), there remains a lack of standardized manufacturing practices for and consensus on appropriate methods and materials for phage production. Current and emerging regulatory guidance discourages the use of animal-derived materials in medicinal product manufacture because of the risk of transmitting adventitious agents (European Medicines Agency 2008, 2025; Food and Drug Administration 2024; MHRA 2025; World Health Organization 2013). This poses a key practical and regulatory question for GMP phage therapy manufacturing: can animal-free culture media be used instead of conventional animal-derived media without compromising phage yield, bactericidal activity, or genomic integrity?

In most research laboratories, phages have been traditionally propagated and studied in bacterial hosts using microbial media containing animal-derived peptones as a source of amino acids for bacterial growth (Atlas 2010; Bonilla and Barr 2018; Gibson et al. 2019). An example of a commonly used animal peptone is tryptone, a digest of bovine milk casein, often prepared with porcine trypsin (Atlas 2010; Gray et al. 2008). Emerging phage therapy guidance documents increasingly advocate that phage products should originate from well-described and validated manufacturing processes (European Medicines Agency 2025; Pirnay et al. 2024a). While some studies have evaluated alternative, animal-free media for scalable phage production (Wiebe et al. 2024), there has been little assessment of how substituting animal-free peptones in place of animal-derived peptones affects phage bactericidal activity, the genomes of phages, and genomes of their propagating hosts. For clinical use, phage preparations must not only achieve sufficiently high titres and retain bactericidal activity, but also demonstrate genomic stability and the absence of undesirable genetic changes throughout manufacturing. Regulatory guidelines highlight whole genome sequencing (WGS) as a key tool for confirming the identity, purity, and safety of phage products (European Medicines Agency 2025; Moon et al. 2025). Yet it is unclear whether switching from animal-derived to animal-free peptones during phage propagation could select for genomic variants in either phages or their bacterial hosts.

Here, we explored whether peptone source of microbial media used for phage propagation affects activity, integrity, and ensuing applicability for animal-free phage manufacture, testing two culture media formulations containing either animal-free soytone or traditional tryptone. We used eight *Pseudomonas aeruginosa* phages and one *Staphylococcus aureus* phage to evaluate whether peptone source affected phage yield and bactericidal activity, hypothesizing that no differences would be observed. In addition to phage phenotypic measures (spot tests, efficiency of plating, and kinetic assays), we performed long-read WGS of paired phage and host bacterial genomes propagated in each medium to test whether animal-free media introduced detectable genomic differences in coding capacity, defense and anti-defense repertoires, or receptor binding proteins compared with traditional media.

## Materials and Methods

### Microbiological Media

Media for bacterial culture and phage propagation are described in Table 1. For ease of interpretation, results assessing bacteria and phages propagated in conventional Luria Bertani media were assigned the abbreviation ‘LB’, while those in animal-free media counterparts were assigned ‘AF’. LB media contained animal-derived tryptone as an amino acid source, while AF media contained animal-free soytone. Broth media for phage propagation were produced and certified according to GMP standards by the manufacturer.

**Table 1.** Microbiological media.

| Media | Manufacturer | Formula | Use |
| --- | --- | --- | --- |
| LB Lennox agar plates | Teknova | 1% tryptone, 0.5% yeast extract, 0.5% NaCl, 1.5% agar | Bacterial culture; phage propagation/ enumeration and EOP |
| LB Lennox agar plates with animal-free soytone | Teknova | 1.69% soytone, 0.5% yeast extract, 0.5% NaCl, 1.5% agar | Bacterial culture; phage propagation/ enumeration |
| LB Lennox agar overlay | Teknova | 1% tryptone, 0.5% yeast extract, 0.5% NaCl, 0.4% agar, 1mM CaCl<sub>2</sub>, 1mM MgCl<sub>2</sub> | Phage propagation (seed stocks for manufacture); phage enumeration, spot test, and EOP |
| LB Lennox agar overlay with animal-free soytone | Teknova | 1.69% soytone, 0.5% yeast extract, 0.5% NaCl, 0.4% agar, 1mM CaCl<sub>2</sub>, 1mM MgCl<sub>2</sub> | Phage propagation (seed stocks for manufacture); phage enumeration |
| LB Lennox broth, GMP manufactured | Teknova | 1% tryptone, 0.5% yeast extract, 0.5% NaCl | Bacterial culture; phage propagation (manufacture for human use) |
| LB Lennox broth with animal-free soytone, GMP manufactured | Teknova | 1.69% soytone, 0.5% yeast extract, 0.5% NaCl | Bacterial culture; phage propagation (manufacture for human use); phage kinetic assay |
| LB Lennox broth | Gibco | 1% tryptone, 0.5% yeast extract, 0.5% NaCl | Phage kinetic assay |

### Bacteria and Phages

*P. aeruginosa* and *S. aureus* clinical isolates used in this study originated from cystic fibrosis lung infections, apart from laboratory reference strain PAO1 (Table 2). Obligately lytic phages targeting these isolates were previously isolated from wastewater samples from the Subiaco Wastewater Plant (Shenton Park, Western Australia, Australia), using a triple plaque purification method (Carr et al. 2025; Iszatt et al. 2022; Ng et al. 2022; Vaitekenas et al. 2022) (Table 2).

**Table 2.**
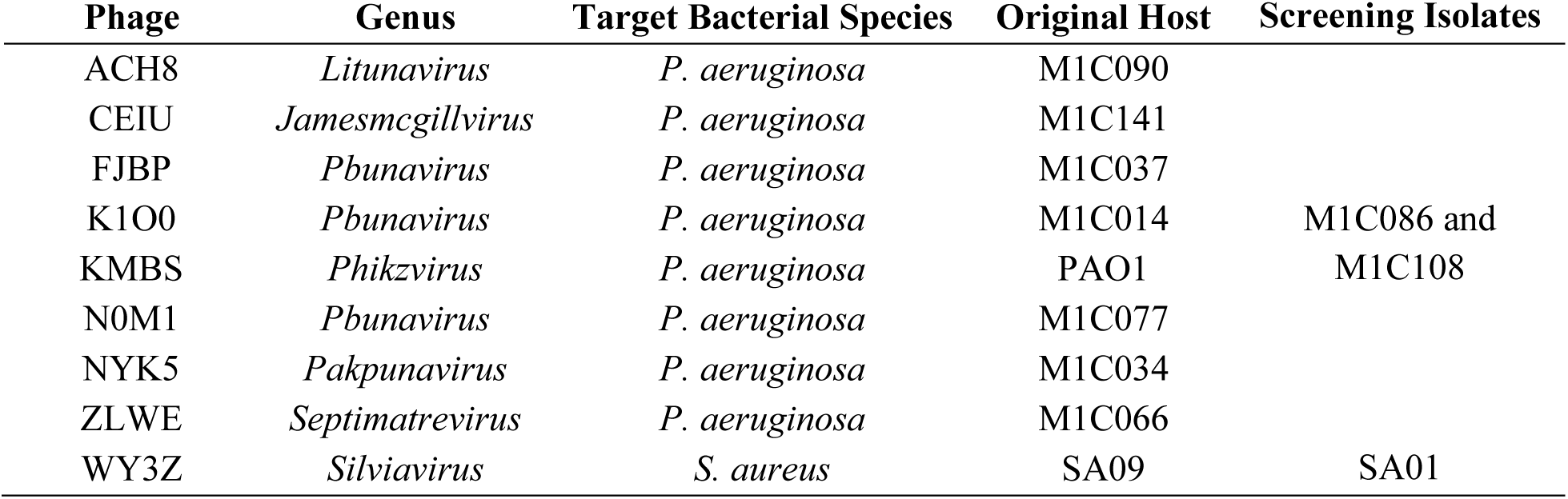
Phages and bacteria.

### Bacterial culture

Both AF and LB bacterial stocks were prepared by culturing isolates in respective broths, mixing with glycerol (25% final concentration), and storing at −80°C. Prior to use in experiments, subcultures of frozen stocks were prepared on AF or LB agar plates, and grown at 37°C for up to 18 hours. A single colony was used to inoculate 5 mL of AF or LB broth and grown up to 18 hours at 37°C at 120rpm.

### Phage Propagation

Phages were propagated to high titers in solid and liquid media (Bonilla and Barr 2018), using both AF or LB formulations. For solid propagation, double agar overlays were prepared by mixing 100 μL of bacterial liquid culture, 100 μL of phage stock, 3 mL of molten overlay agar, and pouring on an agar plate. Once solidified, plates were incubated at 37°C for up to 18 hours, until majority lysis of bacteria was evident. To harvest phages, plates were incubated at room temperature for one hour with 10 mL of phage storage buffer (SM Buffer, gelatine-free; MP Biomedicals), with orbital shaking at 40 rpm. Phage-containing buffer was then collected, centrifuged at 3,000 rpm for 10 minutes to remove debris, filtered via a 0.22 μm membrane to remove residual bacteria, and stored at 4°C until use. For liquid propagation, 10 mL of broth was inoculated with 100 μL of bacterial liquid culture and 100 μL of phage stock, then incubated at 37°C for up to 18 hours at 100 rpm. Phages were similarly harvested by centrifugation to remove debris, then phage-containing supernatants filtered via a 0.22 μm membrane to remove residual bacteria. Filtered supernatants were treated with 1mM CaCl_2_ and 1mM MgCl_2_ for long term phage storage at 4°C.

### Phage Enumeration

Enumeration of infectious phages was performed by plating serial dilutions of phage on bacterial lawns of original propagating hosts (Abedon and Katsaounis 2018; Kropinski et al. 2009). Briefly, a bacterial lawn was prepared via the double agar overlay method by mixing 3 mL of molten overlay agar with 100 μL of bacterial liquid culture. Once solidified, phage dilutions were plated in duplicate 10 μL drops, allowed to dry, and plates incubated at 37°C for up to 18 hours. Small clearings or plaques that formed were counted, and the phage titer calculated [plaque # / (volume factor x dilution factor)] and expressed as plaque forming units per mL (PFU/mL).

### Phage Spot Test

Spot tests were conducted to screen phages for signs of lytic activity against novel bacterial targets as described previously (Kropinski et al. 2009). Briefly, phage stocks were plated in duplicate 10 μL drops on bacterial lawns of naïve bacterial targets, prepared in the same manner as for phage enumeration. After incubation at 37°C for up to 18 hours, spots were visually inspected for lysis and scored as complete lysis, partial lysis, or no lysis.

### Efficiency of Plating

Efficiency of Plating (EOP) was used to measure phage virulence against novel bacterial targets as previously described (Abedon and Katsaounis 2018; Kropinski et al. 2009). Similar to the process for titrating phages, serial dilutions of phages were plated in duplicate drops on bacterial lawns of original propagating hosts and a novel bacterial target. After incubation at 37°C for up to 18 hours, phage titers were calculated, and EOP scores determined by dividing phage titer in a novel target by the phage titer in its original propagating host. These EOP values were then used to categorize phages as highly virulent (0.1<EOP>1), moderately virulent (0.001<EOP>0.1), avirulent but active (EOP<0.001), or avirulent (EOP=0), based on a published metric (Gibson et al. 2019).

### Kinetic Assays

Kinetic assays were used to assess phage suppression of bacterial growth over time via turbidity measurements of liquid cultures at OD600nm (Rajnovic et al. 2019). Briefly, on a 96 well plate, 10^6^ bacterial colony forming units (CFU) and 10^6^ phage PFU (multiplicity of infection [MOI] of one) were combined in 200 µL of broth per well. Wells with bacteria alone served as positive controls for bacterial growth, wells with phage alone served as negative controls for bacterial growth, and wells with broth alone served as blanks. Assay plates were set in a Spark Microplate Reader (Tecan) and incubated for 16 hours at the following parameters (37°C, OD600nm reads every 10 minutes, with 96rpm orbital shaking in between reads). Values were blank corrected then plotted as growth curves over time. Curves were assigned kinetic forms based on published metrics from the Israeli Phage Therapy Centre (Yerushalmy et al. 2023). A Sensitive result was characterized by an almost fully flat curve with low signal throughout the assay period, indicative of sustained bacterial inhibition. A Sensitive + regrow result was characterized by a flat curve followed by bacterial regrowth after more than 5 hours. An Intermediate result was characterized by some observable bacterial inhibition and/or late lysis or stationary growth. A Resistant result was characterized by little to no difference in bacterial growth with or without phage present. Kinetic assay results were quantified via area under the curve (AUC) analysis, which was used to calculate the suppression index for each phage [(AUC_no phage_ – AUC_phage_) / AUC_no phage_)] (Kim et al. 2024).

### DNA Extraction

Bacterial and phage genomic DNA were extracted and purified using the DNeasy Blood and Tissue Kit (Qiagen) as previously described (Hedges 2025; Mitchell and Samuel 2025). Briefly, for phage DNA extractions an aliquot of phage lysate was treated with 5 μL of both DNase I and RNase A for 30 minutes at 37°C to remove free bacterial host nucleic acids, then treated with 180 μL of tissue lysis buffer (Buffer ATL) and 20 μL of proteinase K for 15 minutes at 56°C to release phage DNA. For Gram-negative bacterial DNA extraction, cell pellets were washed with 0.1M EDTA, after which Buffer ATL and proteinase K were added to lyse bacterial cells and release DNA. Any contaminating RNA was then removed by treatment with 5 μL of RNAse A for 15 minutes at 37°C. For Gram-positive bacterial DNA extraction, cell pellets were first washed with 0.1M EDTA then washed again in 1X phosphate buffered saline (PBS), after which 2,500 units/sample of metapolyzyme in 100 μL of PBS was added to promote cell lysis and DNA release, followed by a subsequent RNAse A treatment. Liberated DNA was purified on spin columns per manufacturer’s instructions (Qiagen). DNA concentrations were measured using a Qubit Flex Fluorometer and the Quant-iT 1× dsDNA Assay Kit, with the Broad Range kit used for bacterial DNA and the High Sensitivity kit for phage DNA (ThermoFisher). DNA integrity was assessed by running samples on a 2% agarose gel and visualising bands following DNA staining.

### Whole Genome Sequencing

DNA was neither sheared nor size selected before library preparation. Long-read WGS of bacterial and phage DNA was performed using the Oxford Nanopore PromethION 2 Solo. Libraries were prepared using either the Native Barcoding Kit 24 V14 (SQK-NBD114.24; Oxford Nanopore Technologies, UK) or Rapid PCR Barcoding Kit 24 V14 (SQK-RPB114.24; Oxford Nanopore Technologies, UK) and sequenced using PromethION 10.4.1 flow cells (FLO-PRO114M). Raw reads from phages were basecalled and demultiplexed using Dorado within the MinKNOW software (v24.06.5) using the dna_r10.4.1_e8.2_400bps_sup@v4.3.0 basecalling model, while raw reads from bacteria were basecalled and demultiplexed using stand-alone dorado (v1.0.0) and the dna_r10.4.1_e8.2_400bps_sup@v5.2.0 basecalling model.

### Bioinformatics

For bacterial WGS, genome assembly and annotation were performed using a publicly available workflow (www.github.com/samuelmontgomery/bacass) (Montgomery 2025). Briefly, raw reads were length and quality filtered (>Q10, >1000bp) using chopper (v0.10.0) (De Coster and Rademakers 2023), checked for contamination using kraken2 (v2.1.5) (Wood et al. 2019) with the standard 16GB database, and visualised using NanoPlot (v1.44.1) (De Coster and Rademakers 2023). Bacterial genomes were assembled from quality-controlled reads using flye (v2.9.6) (Kolmogorov et al. 2019), a consensus assembly generated using autocycler (v0.5.2), and genome quality was assessed using minimap2 (v2.29) (Li 2018), samtools (v1.22.0) (Li et al. 2009), qualimap (v2.3) (Garcia-Alcalde et al. 2012) and checkm (v1.2.3) (Parks et al. 2015). The resulting bacterial genomes were annotated using bakta (v1.11.0) (Schwengers et al. 2021) using the full database (v6.0) with further specialist annotation of bacterial defense systems performed with PADLOC and its database (v2.0.0) (Payne et al. 2021). Prophage sequences integrated into bacterial genomes were identified using genomad (v1.11.0) (Camargo et al. 2024). Differences in bacterial genomes between the two growth mediums were assessed by variant calling using clair3 (v1.1.1) (Zheng et al. 2022). For phage WGS, raw reads were length and quality filtered (>Q10, >1000bp) using chopper (v0.10.0) (De Coster and Rademakers 2023), subsampled to 500X coverage with rasusa (v2.1.0) (Hall 2022) and visualized using NanoPlot (v1.44.1) (De Coster and Rademakers 2023). Genome assembly was conducted using flye (v2.9.6) (Kolmogorov et al. 2019) and quality of the resulting genomes were assessed using minimap2 (v2.29) (Li 2018), qualimap (v2.3) (Garcia-Alcalde et al. 2012) and checkv (v1.0.3) (Nayfach et al. 2021). Phage genomes were then annotated using Pharokka (v1.7.5) (Bouras et al. 2023) and the PHROGS database (Terzian et al. 2021) followed by specialist annotation of receptor binding proteins (RBPs) and anti-defenses performed using PhageRBPdetection v3.0.0 (Boeckaerts et al. 2022) and dbAPIs (commit ID: 74af7e2) (Yan et al. 2024). Average nucleotide identity (ANI) between the two assemblies was calculated using ANI calculator (Yoon et al. 2017). Genomic differences between phages propagated in the two growth media were evaluated through variant calling using clair3 (v.1.0.10) (Zheng et al. 2022), using the LB assembly as the reference.

### Statistical Analysis

Statistical analyses, AUC data, and figures were prepared using GraphPad Prism v10 (GraphPad Software, La Jolla, CA, USA). Phage propagations and subsequent tests and WGS were performed using eight different phages targeting *P. aeruginosa* and one phage targeting *S. aureus*, encompassing nine biological replicates. Phage screening results were obtained using two *P. aeruginosa* isolates and one *S. aureus* isolate, encompassing three biological replicates for screening assays. Data from stock propagations, kinetic assays, spot tests, and efficiency of plating are shown as averages from two technical replicates. To assess differences in outcomes between AF and LB media, propagated phage titers and phage suppression indices were compared via multiple Mann-Whitney tests, and comparisons of AUC results were performed via a two-way ANOVA. For all statistical tests a Bonferroni correction for multiple comparisons was applied, with p<0.05 considered statistically significant.

## Results

### Phage propagation

To determine if media peptone source affected phage yield during stock propagation, eight phages targeting *P. aeruginosa* and one phage targeting *S. aureus* were propagated in solid and liquid media. All phages grew to high titers of ≥10^8^ PFU/mL across both propagation methods, with no significant differences in phage yield following propagation in either AF or LB base media (Figure 1). Two exceptions include solid propagation of phage WY3Z and liquid propagation of CEIU, which showed >1 log_10_ increases in titer when propagated in AF media (Figure 1). Altogether these results indicate peptone source is not a strong determinant of phage yield, with either traditional or animal-free microbiological media suitable for high titer phage propagation.

**Figure 1.**
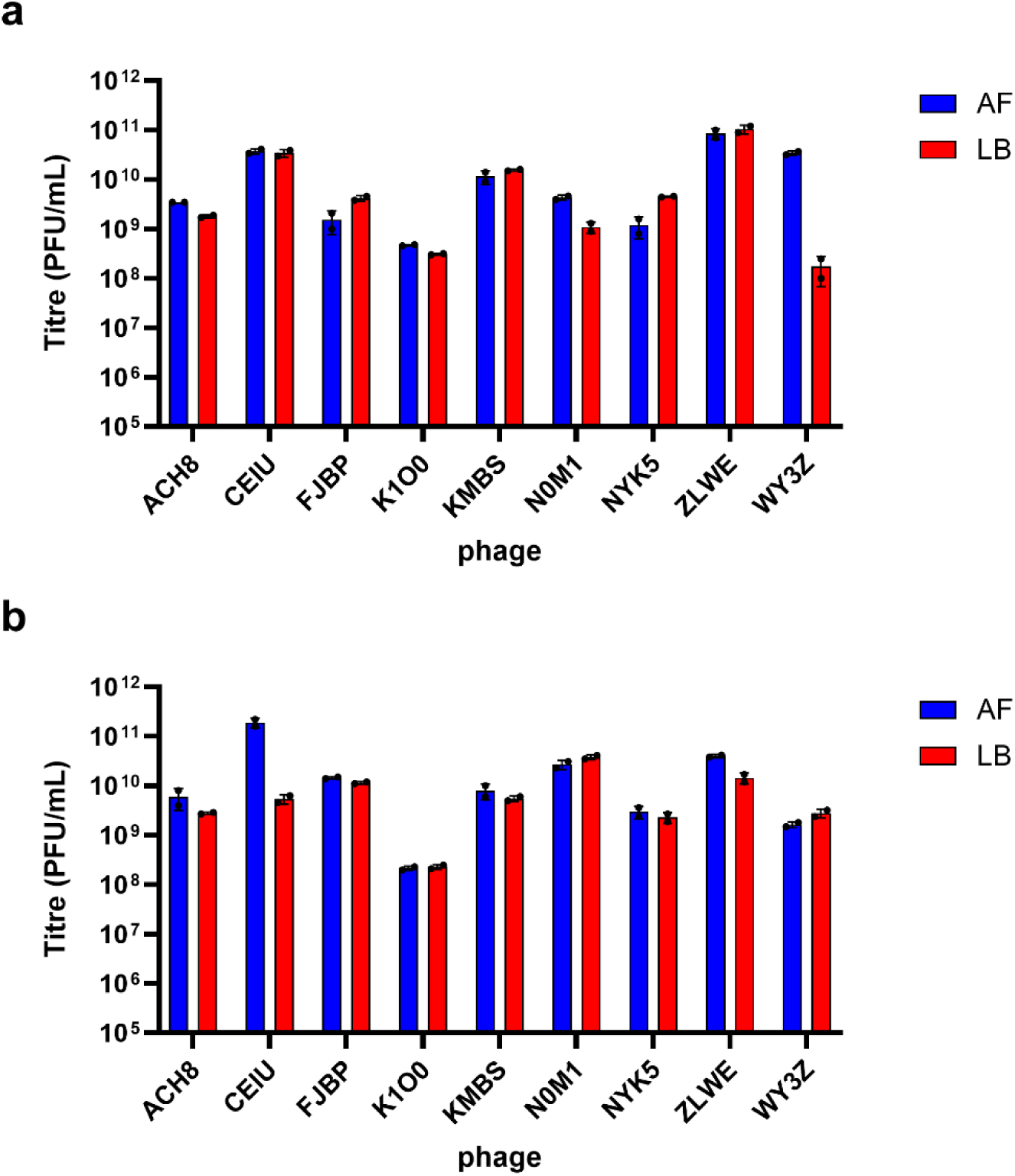
Phage propagation with different peptone sources. Phage yields (PFU/mL) of a) agar-propagated and b) broth-propagated phages in both LB and AF media formulations. Data is shown as the mean ± SD of two technical replicates.

### Suppression of bacterial growth

Kinetic assays were performed to assess if peptone source used for stock propagation affected phage suppression of bacterial growth. Notably, bacterial growth in positive control wells typically reached higher turbidity at stationary phase, and for some isolates earlier log phase growth, when incubated in AF versus LB broth (Figures 2 and 3). Despite differences across positive controls for bacterial growth, scoring of phage-induced bacterial suppression was the same for all phages regardless of peptone source, and results were consistent whether the phage stock was propagated using a solid overlay or broth (Figures 2 and 3; Supplementary Table 1). Kinetic assay results of both agar and broth-propagated phages were combined for AUC analysis, which similarly showed significant increases in AUC values for bacterial hosts growing in AF versus LB broth (p<0.001), but no significant differences in AUC when phage were present (Figure 4a). Furthermore, there were no significant differences in phage suppression indices between AF and LB groups (Figure 4b).

**Figure 2.**
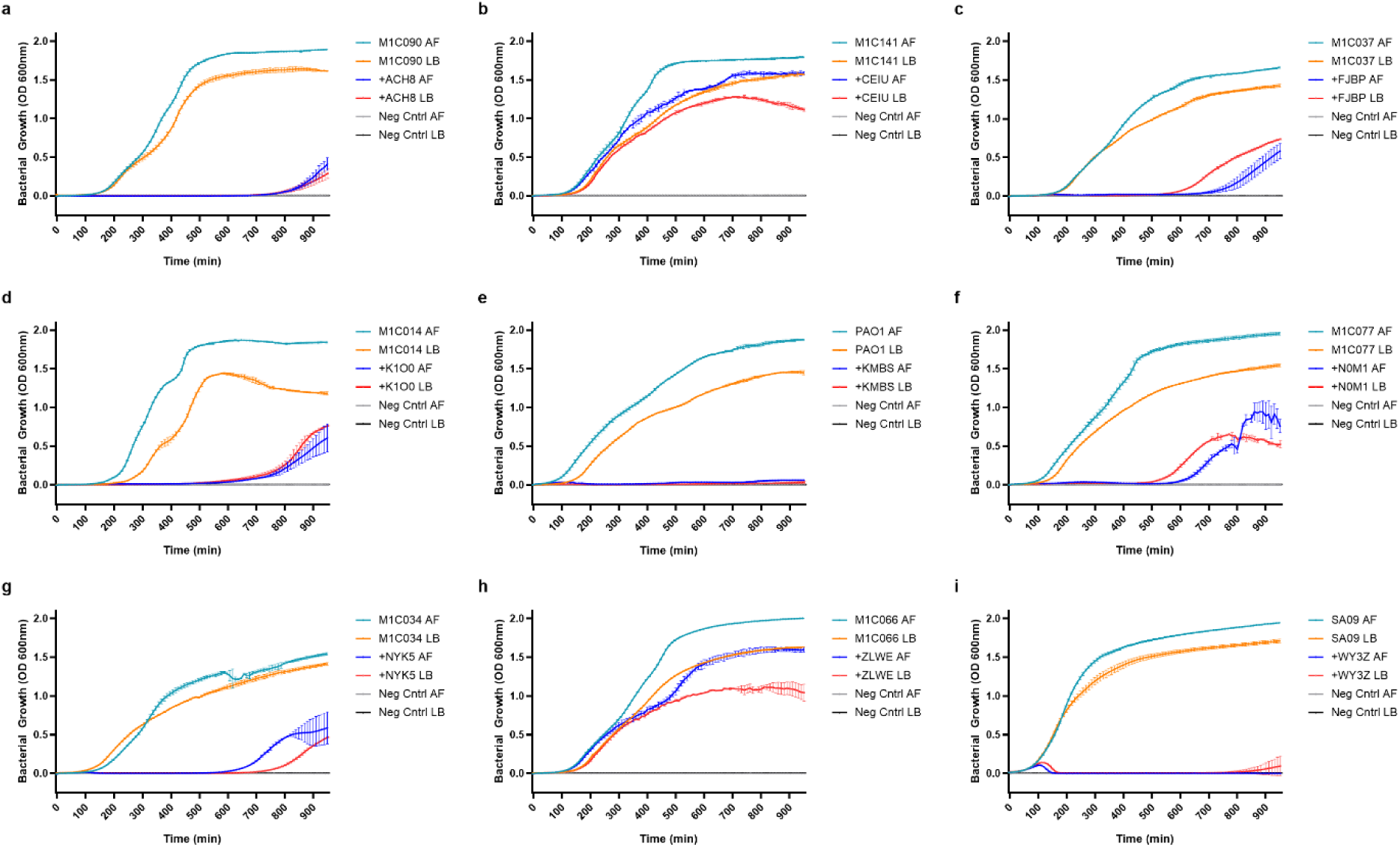
Peptone source of solid propagation media does not inhibit phage suppression of bacterial growth. Growth curves from kinetic assays testing agar-propagated phages a) ACH8, b) CEIU, c) FJBP, d) K1O0, e) KMBS, f) N0M1, g) NYK5, h) ZLWE and i) WY3Z. Data is shown as the mean ± SD of two technical replicates.

**Figure 3.**
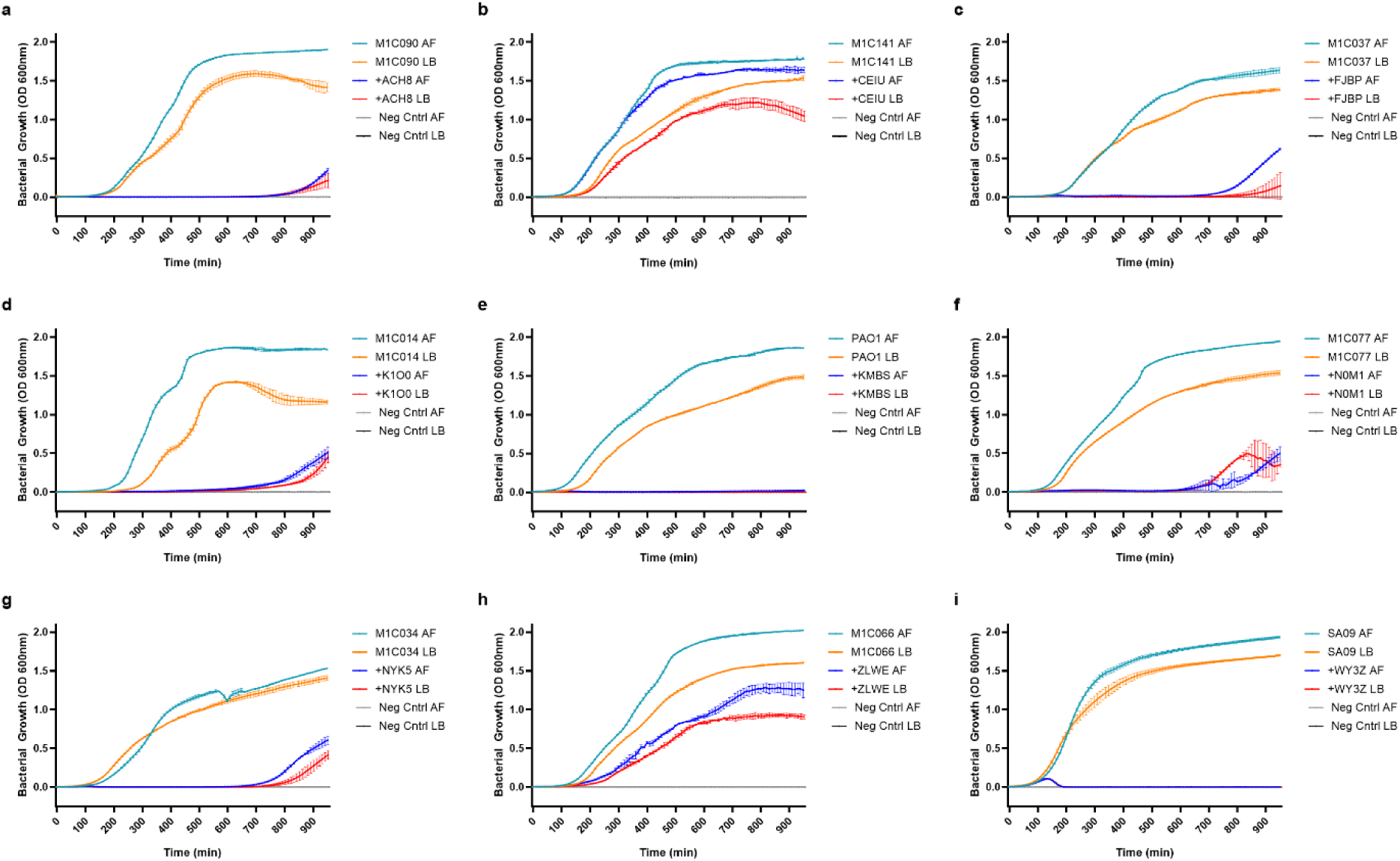
Peptone source of liquid propagation media does not inhibit phage suppression of bacterial growth. Growth curves from kinetic assays testing broth-propagated phages a) ACH8, b) CEIU, c) FJBP, d) K1O0, e) KMBS, f) N0M1, g) NYK5, h) ZLWE and i) WY3Z. Data is shown as the mean ± SD of two technical replicates.

**Figure 4.**
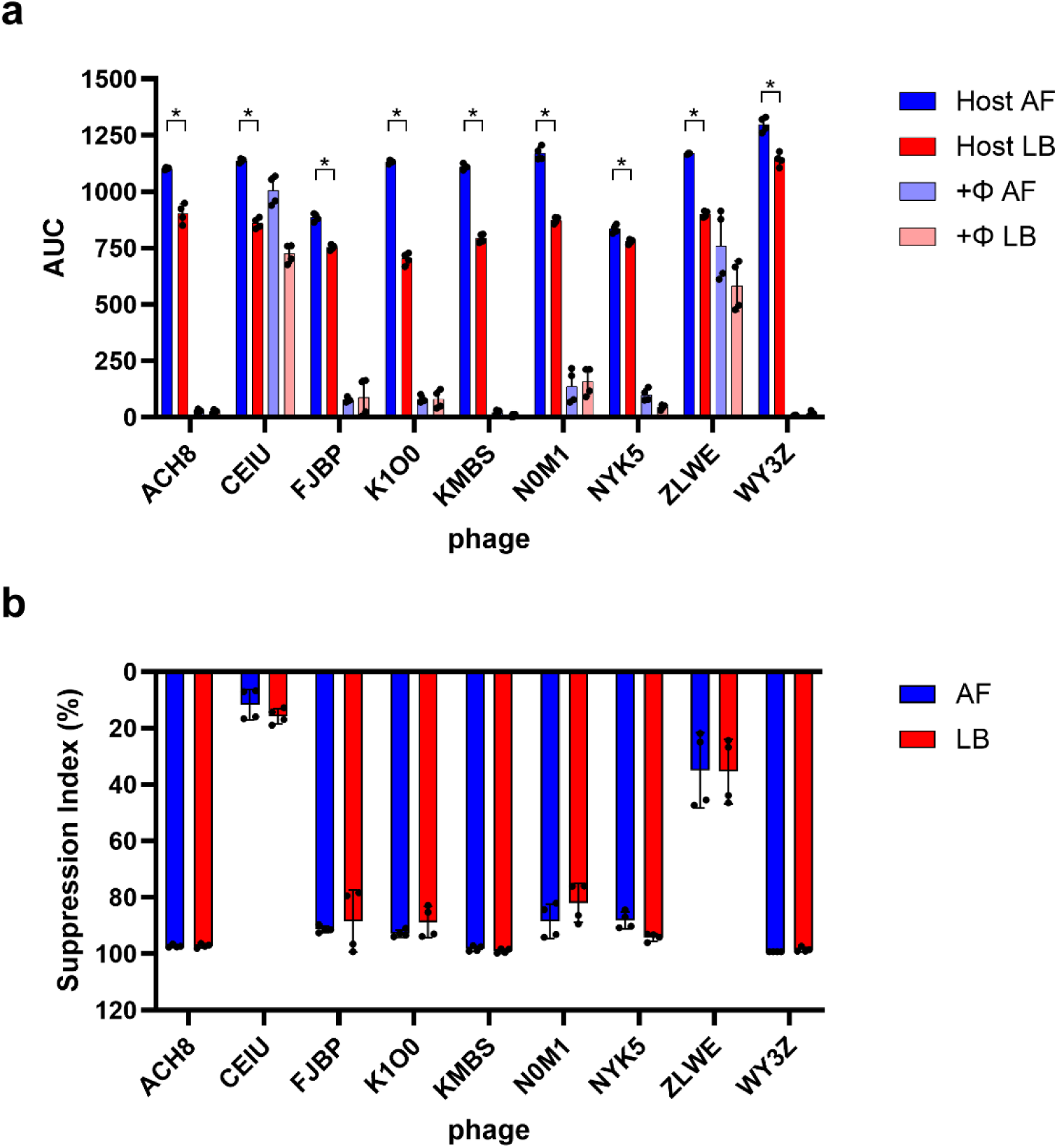
Peptone source alters bacterial growth but not phage suppression. a) Bacterial hosts had significantly higher AUC values when growing in AF versus LB broth (*p<0.001, two-way ANOVA with a Bonferroni correction). Despite differences in bacterial growth in the absence of phages, no significant differences were observed among b) phage suppression indices. Data is shown as mean ± SD of combined technical replicates from solid and liquid propagated phages (n=4).

### Phage screening tests

To determine if peptone source affected phage screening results, mock screens were then conducted against two additional *P. aeruginosa* isolates (M1C086 and M1C108) and an additional *S. aureus* isolate (SA01). Screens were conducted using phages propagated in solid media from both AF and LB formulations, with assays run using LB plates and broth. Spot test scores were identical between phages raised in AF and LB media (Figure 5a). Following this EOP assays were performed using phage ACH8 since it showed lytic activity against both *P. aeruginosa* test isolates in the spot test. *Staphylococcus* phage WY3Z was also tested by EOP, despite being lysis-negative in the spot test, as a control for phage resistance. EOP values against each isolate did vary approximately two-fold between AF and LB-raised phages; however, the resulting virulence score was identical for each phage, showing no difference in the interpretation of these results between peptone sources (Table 3). These two phages were then also tested in kinetic assays against their respective test isolates, as well as original propagating hosts. Once again, patterns of bacterial growth suppression observed in kinetic forms did not differ between AF and LB-raised phages, and results were consistent for growth curves showing phage sensitivity or resistance (Figure 5b-f and Supplementary Table 2).

**Figure 5.**
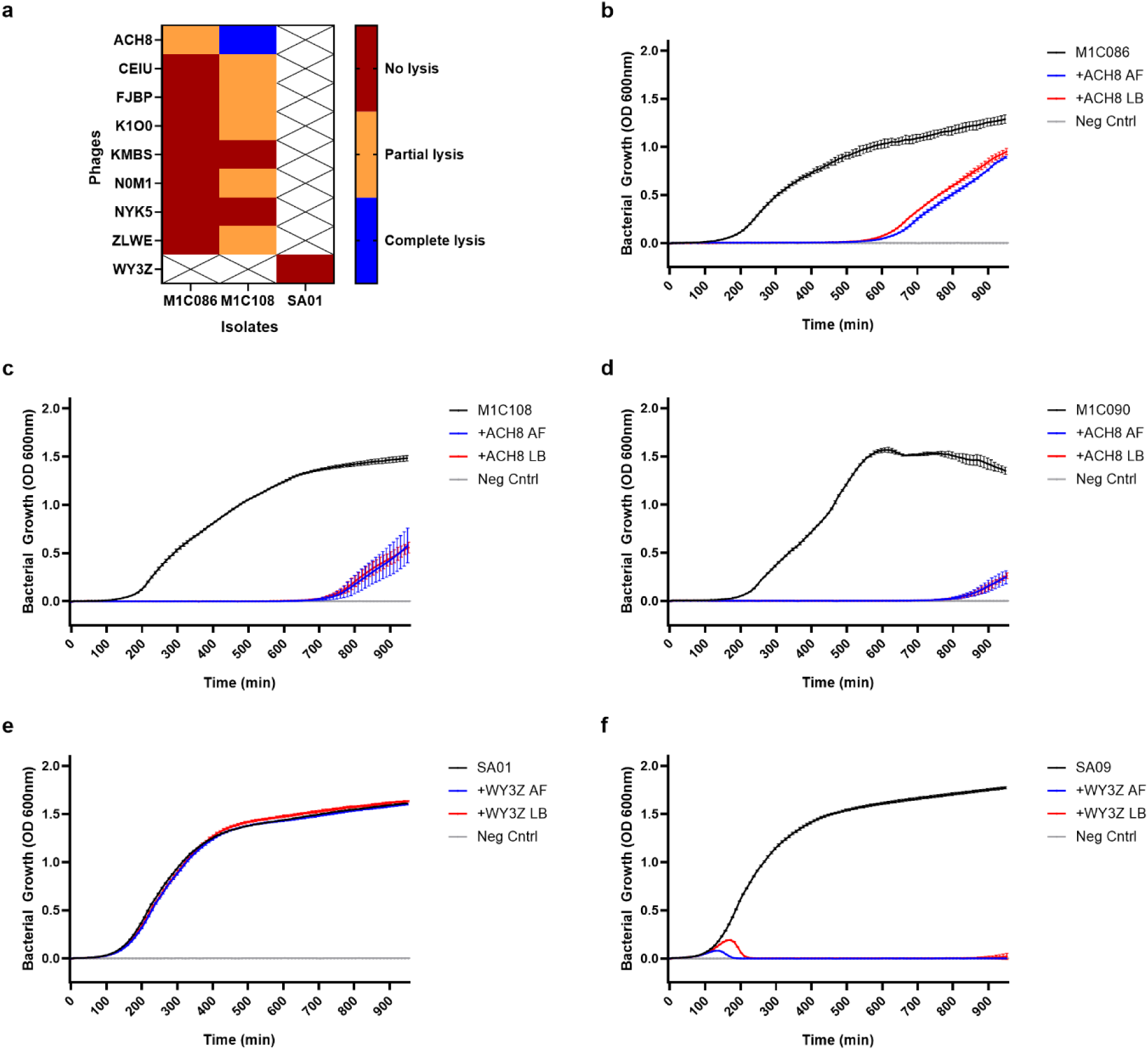
Phage screening results are unaffected by peptone source used for stock propagation. a) Spot test results of phages on novel bacterial targets. Kinetic assay results for phage ACH8 tested on *P. aeruginosa* isolates b) M1C086, c) M1C108, and original host d) M1C090; results for phage WY3Z tested on *S. aureus* isolate e) SA01 and original host f) SA09. Data is shown as the mean ± SD of two technical replicates.

**Table 3.** Peptone source of phage propagation media does not alter EOP scores. (LB=Luria Bertani, AF=animal-free, n/a=not applicable, EOP=efficiency of plating)

| Phage | Media Type | Isolate | PFU/mL | EOP | Virulence Score |
| --- | --- | --- | --- | --- | --- |
| ACH8 | AF | Original host | 4.65 x 10<sup>9</sup> | 1.00 | n/a |
|  | LB | Original host | 1.80 x 10<sup>9</sup> | 1.00 | n/a |
|  | AF | M1C086 | 6.65 x 10<sup>8</sup> | 0.143 | Highly Virulent |
|  | LB | M1C086 | 4.45 x 10<sup>8</sup> | 0.247 | Highly Virulent |
|  | AF | M1C108 | 3.40 x 10<sup>9</sup> | 0.731 | Highly Virulent |
|  | LB | M1C108 | 2.80 x 10<sup>9</sup> | 1.556 | Highly Virulent |
| WY3Z | AF | Original host | 2.20 x 10<sup>10</sup> | 1.00 | n/a |
|  | LB | Original host | 1.60 x 10<sup>8</sup> | 1.00 | n/a |
|  | AF | SA01 | 0 | 0 | Avirulent |
|  | LB | SA01 | 0 | 0 | Avirulent |

### Whole genome sequencing of phages and bacterial hosts

To assess any genomic differences arising from propagation in either AF or LB media, we performed WGS on bacterial isolates and phages grown in broths of both formulations. All bacterial and phage genomes were successfully assembled and annotated (Supplementary Tables 3 and 4, respectively). There were no significant differences in assembled bacterial genome size or predicted gene annotations (Table 4), with a small number of variants (single nucleotide polymorphisms and insertions/deletions) identified between AF and LB broth (Supplementary Table 5). Additionally, there were no differences in the number or type of defense systems annotated in the bacterial isolates grown in either the AF or LB broth (Supplementary Table 6). No substantial differences were observed in the assembled genome size or predicted gene annotations between phages propagated in AF and LB broth, and ANI (%) between these genomes was found to be >99% for all phages (Table 5). Only 5 variants were identified between AF and LB-grown phages, primarily within hypothetical proteins and one virion structural protein (Supplementary Table 7); however, the number and types of anti-defenses and receptor binding proteins (RBPs) were consistent across phages grown in both formulations (Figure 6). Importantly, phages propagated in AF media remained obligately lytic as did their LB phage counterparts, with an absence of integrase genes and other markers of lysogeny. Furthermore, both AF and LB phages did not transduce any deleterious bacterial genes, including genes associated with increased bacterial virulence or AMR.

**Figure 6.**
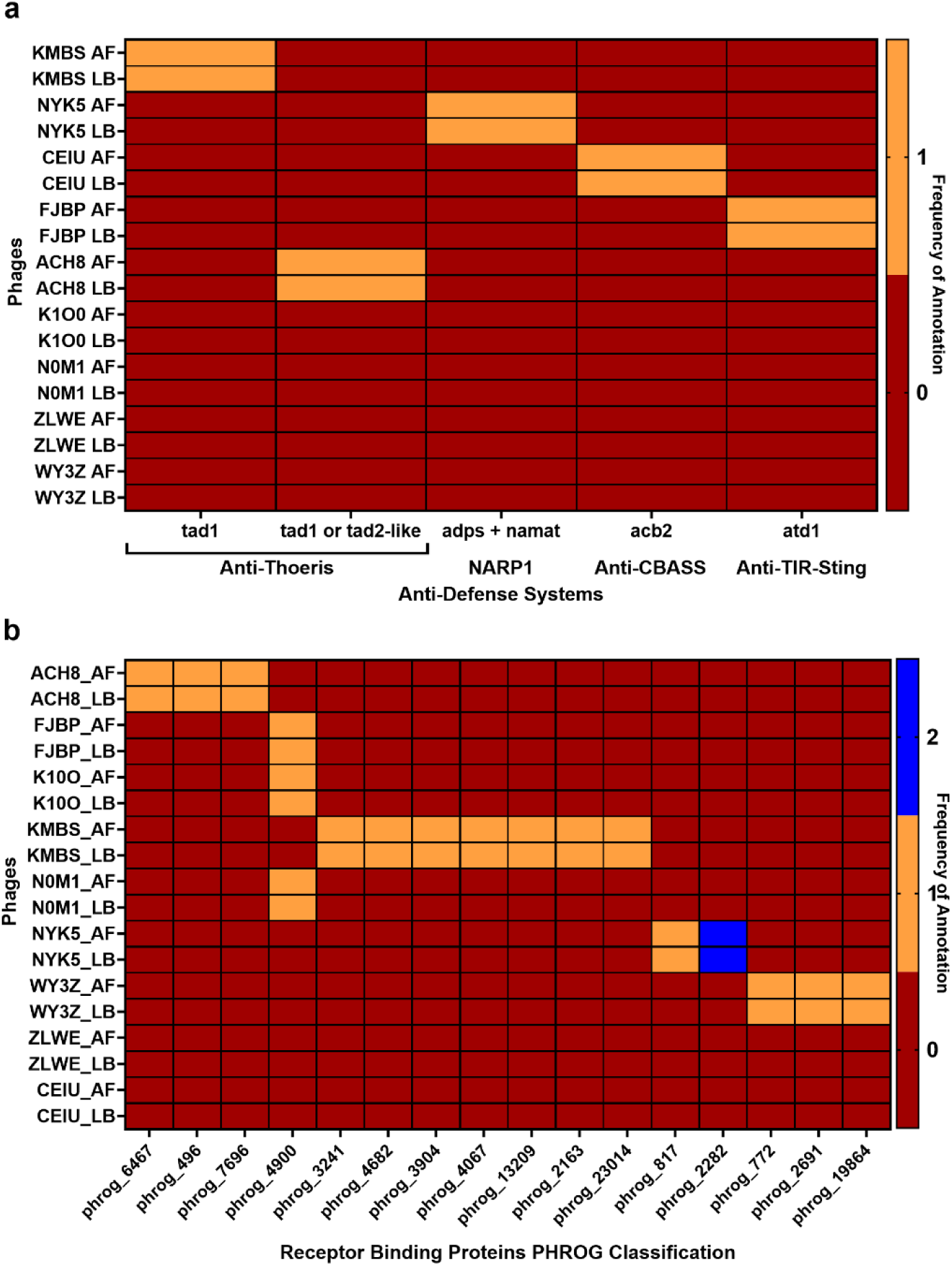
Annotated phage anti-defense and phage receptor binding protein profiles. Colors indicate the frequency of a) phage anti-defenses and b) receptor binding proteins identified in phage genomes. Receptor binding proteins have been classified according to their Prokaryotic Virus Remote Homologous Groups (PHROG) number.

**Table 4.** Peptone source does not significantly alter bacterial genomes or proteins.

| Isolate | Media type | Genome length (bp) | Contigs | tRNAs | ncRNAs | CRISPR arrays | CDSs |
| --- | --- | --- | --- | --- | --- | --- | --- |
| M1C014 | AF | 6601387 | 1 | 66 | 47 | 0 | 5995 |
|  | LB | 6601386 | 1 | 66 | 47 | 0 | 5997 |
| M1C034 | AF | 6563285 | 1 | 64 | 44 | 3 | 5977 |
|  | LB | 6563285 | 1 | 64 | 44 | 3 | 5977 |
| M1C037 | AF | 6618444 | 1 | 65 | 47 | 0 | 5992 |
|  | LB | 6618444 | 1 | 65 | 47 | 0 | 5992 |
| M1C054 | AF | 6329616 | 1 | 63 | 47 | 4 | 5737 |
|  | LB | 6329616 | 1 | 63 | 47 | 4 | 5731 |
| M1C066 | AF | 6400394 | 1 | 66 | 47 | 0 | 5819 |
|  | LB | 6400393 | 1 | 66 | 47 | 0 | 5816 |
| M1C077 | AF | 6362869 | 1 | 63 | 45 | 0 | 5772 |
|  | LB | 6362864 | 1 | 63 | 45 | 0 | 5773 |
| M1C090 | AF | 6230865 | 1 | 65 | 47 | 0 | 5647 |
|  | LB | 6230865 | 1 | 65 | 47 | 0 | 5647 |
| M1C093 | AF | 6354068 | 1 | 64 | 48 | 3 | 5794 |
|  | LB | 6354068 | 1 | 64 | 48 | 3 | 5794 |
| M1C141 | AF | 6832133 | 1 | 69 | 54 | 2 | 6334 |
|  | LB | 6832133 | 1 | 69 | 54 | 2 | 6334 |
| PA01 | AF | 6263669 | 1 | 63 | 48 | 0 | 5680 |
|  | LB | 6263669 | 1 | 63 | 48 | 0 | 5680 |
| SA01 | AF | 2811201 | 1 | 59 | 76 | 0 | 2601 |
|  | LB | 2811201 | 1 | 59 | 76 | 0 | 2601 |
| SA09 | AF | 2844927 | 2 | 61 | 81 | 0 | 2599 |
|  | LB | 2844927 | 2 | 61 | 81 | 0 | 2599 |

**Table 5.** Peptone source does not significantly alter phage genomes or proteins.

| Phage | Media type | Genus | Genome length (bp) | GC% | CDSs | ANI% |
| --- | --- | --- | --- | --- | --- | --- |
| ACH8 | AF | *Litunavirus* | 72936 | 54.75 | 101 | 100 |
|  | LB |  | 72945 | 54.75 | 101 |  |
| CEIU | AF | *Jamesmcgillvirus* | 43225 | 45.06 | 56 | 100 |
|  | LB |  | 43225 | 45.06 | 56 |  |
| FJBP | AF | *Pbunavirus* | 66524 | 55.56 | 100 | 100 |
|  | LB |  | 66524 | 55.56 | 100 |  |
| K100 | AF | *Pbunavirus* | 66464 | 55.57 | 108 | 100 |
|  | LB |  | 66464 | 55.57 | 108 |  |
| KMBS | AF | *Phikzvirus* | 277066 | 36.81 | 405 | 100 |
|  | LB |  | 277066 | 36.81 | 405 |  |
| N0M1 | AF | *Pbunavirus* | 66157 | 55.68 | 100 | 100 |
|  | LB |  | 66157 | 55.68 | 100 |  |
| NYK5 | AF | *Pakpunavirus* | 92692 | 49.39 | 193 | 99.99 |
|  | LB |  | 92696 | 49.39 | 193 |  |
| WY3Z | AF | *Silviavirus* | 135360 | 30.09 | 218 | 99.88 |
|  | LB |  | 135360 | 30.09 | 215 |  |
| ZLWE | AF | *Septimatrevirus* | 43066 | 54.53 | 59 | 100 |
|  | LB |  | 43067 | 54.53 | 60 |  |

## Discussion

We observed similar phage yield and bactericidal activity across phages propagated with either animal-free soytone or tryptone as an amino acid source, and no substantial genomic differences in phages and bacterial hosts grown in AF or LB media. These findings were consistent across the different phages we tested, confirming our hypothesis that media peptone sources do not result in significant differences in propagated phages. Our findings reflect results from a previous investigation of scalable production of phages targeting *Escherichia coli*, in which phage growth and activity were compared in LB Lennox media versus animal-free Select APS media, which uses a soy hydrolysate in place of tryptone (Wiebe et al. 2024). Phage yields, bactericidal activity, and bacterial growth were similar in both media types, as was scalability of phage production and targeting of novel hosts (Wiebe et al. 2024). Together with our observations, this suggests that AF media performs as well as LB media for propagation of phages for therapeutic use. Furthermore, we did not observe strong differences in genomes of phages and their hosts according to the peptone source used for propagation. Notably, phages propagated in AF media retained desirable genetic characteristics for therapeutic use identical to LB-propagated phages, chiefly a lack of lysogenic markers or harmful bacterial genes (European Medicines Agency 2025; Gibson et al. 2019). Verifying genomic stability of both phages and bacterial hosts used in production will be crucial for manufacture of medicinal phages. Genomic characterization is a key metric of identity and safety of the final phage product, with WGS mentioned in current and emerging phage therapy guidelines (European Medicines Agency 2025; Moon et al. 2025).

Here, we also tested two phage propagation methods using solid and liquid media. Solid propagation in agar is a common method used in research laboratories, with phages harvested in a stabilising buffer suitable for long term phage storage (Bonilla and Barr 2018). Liquid propagation in broth is also used in research settings, but can require supplementation with salts to maintain phage stability (Bonilla and Barr 2018). We tested both propagation methods, with broth representative of the method that is scalable for large scale propagation (Rebula et al. 2023; Wiebe et al. 2024). Both methods yielded high phage titers, with no difference in kinetic forms produced by phages propagated using either method, suggesting propagation method does not influence phage bactericidal activity.

In addition to phage propagation, our findings provide insights on media for *in vitro* phage susceptibility assays, as there is no consensus on the appropriate medium for clinical testing (Kolenda et al. 2024; Yerushalmy et al. 2023). Studies note that the choice of media used for phage assays can have an effect on plaque formation (Ramesh et al. 2019) or bacterial growth rate and phage activity in liquid broth (Nabergoj et al. 2018; Wiebe et al. 2024). While we did not assess plaque morphology in our study, no reduction in countable plaques was evident when using AF vs LB overlay, suggesting the animal-free alternative did not inhibit plaque formation and consequently impair phage enumeration. Additionally, in a mock therapeutic phage screen, we observed no differences in assay outcomes when phages raised in either LB or AF media were tested on both solid and liquid LB media. This implies that phages raised in AF medium can be tested in LB media, with no indication that this would compromise measures of bactericidal activity.

To our knowledge, this study is the first to purposefully assess effects of animal-free peptone on phage activity and genomic integrity in the context of therapeutic production. While we tested culture media formulations that are relevant in manufacturing settings, this study was limited by use of small-scale phage propagation methods instead of large-scale processes (Wiebe et al. 2024). In addition, although we did not see genomic differences in phage or bacteria between media formulations, dual RNA-seq may provide further insights into host-pathogen dynamics and potential peptone-associated differences in phage and bacterial gene expression (Putzeys et al. 2023; Wicke et al. 2021). Lastly, not all bacterial pathogens, and consequently phages targeting them, can be cultured with animal-free media (Lagier et al. 2015). Therefore, our findings may not be applicable to fastidious bacteria that require animal-derived reagents for growth in culture. In this case, manufacturers of phage therapy products targeting these microbes will need to justify use of animal-derived reagents to regulators and conduct appropriate risk analyses and sourcing of materials in use to reduce potential hazards for patients.

In summary, we have shown that an animal-free peptone source in culture media does not alter phages propagated for therapeutic use. Findings support the use of animal-free media for phage production where animal-derived ingredients are not desired, such as in GMP manufacture of therapeutic goods. Enabling medicinal manufacture of phage products is an important step to ensure safety and quality for patients, reinforce compliance with regulatory guidelines, support large scale clinical trials, and facilitate full translation of phage therapy.

## Acknowledgements

We thank Professor Barbara Chang at the University of Western Australia for providing *P. aeruginosa* reference strain PAO1. We thank Prof. Sarath Ranganathan and Ms. Rosemary Carzino from the Murdoch Children’s Research Institute (MCRI) for providing the *P. aeruginosa* clinical isolates used in this study. We thank Professor Scott Bell at Griffith University for providing *S. aureus* isolates SA01 and SA09. We also acknowledge that this project was conducted on the traditional homelands of the Noongar people, with phages isolated from waters across Noongar Wadjak. Phage WA thanks the Sharon Gregory and Walyalap Waangkan Noongar language team, who named one of the phages used in this study in the Wadjak Noongar language. Karil-mokiny-kep-djiraly-karakaata-Wadjak-1 (phage WY3Z), abbreviated as Karil-mokiny-1, translates as “crab-like (from) water northern karrakatta-wadjak.”

## Author Contributions

D.R.L. (Conceptualization, Investigation, Formal Analysis, Writing-original draft), P.G.C. (Investigation, Formal Analysis, Writing-original draft), M.G.H (Investigation, Formal Analysis, Writing-original draft), A.V. (Investigation, Formal Analysis, Writing-original draft), Z.V. (Funding Acquisition, Supervision, Writing-review and editing), S.M.S. (Funding Acquisition, Writing-review and editing), S.T.M. (Investigation, Formal Analysis, Writing-original draft), A.K. (Funding Acquisition, Investigation, Supervision, Writing-review and editing).

## Data Availability

The raw sequencing reads and assembled genomes generated in this study have been deposited in the NCBI database under BioProject accession numbers PRJNA862674 (*Pseudomonas aeruginosa*), PRJNA862679 (*Staphylococcus aureus*), and PRJNA862681 (bacteriophages). These data are publicly available through the NCBI Sequence Read Archive (SRA) and GenBank under the corresponding BioProject identifiers. Supplementary data are available in the online supplementary material.

## Abbreviations

AF: Animal-free
LB: Luria Bertani
CFU: colony forming units
PFU: plaque forming units
EOP: Efficiency of Plating
GMP: Good Manufacturing Practice

