## Supplementary Table for "Animal-free peptones do not alter bacteriophages propagated for therapeutic use": Supplementary Tables 1-4 & 7.pdf

| Phage | Propagation Method | Media Type | Kinetic Form |
|---|---|---|---|
| ACH8 | Solid | AF | Sensitive + regrow |
| | | LB | Sensitive + regrow |
| | Liquid | AF | Sensitive + regrow |
| | | LB | Sensitive + regrow |
| CEIU | Solid | AF | Resistant |
| | | LB | Resistant |
| | Liquid | AF | Resistant |
| | | LB | Resistant |
| FJBP | Solid | AF | Sensitive + regrow |
| | | LB | Sensitive + regrow |
| | Liquid | AF | Sensitive + regrow |
| | | LB | Sensitive + regrow |
| K1O0 | Solid | AF | Sensitive + regrow |
| | | LB | Sensitive + regrow |
| | Liquid | AF | Sensitive + regrow |
| | | LB | Sensitive + regrow |
| KMBS | Solid | AF | Sensitive |
| | | LB | Sensitive |
| | Liquid | AF | Sensitive |
| | | LB | Sensitive |
| N0M1 | Solid | AF | Sensitive + regrow |
| | | LB | Sensitive + regrow |
| | Liquid | AF | Sensitive + regrow |
| | | LB | Sensitive + regrow |
| NYK5 | Solid | AF | Sensitive + regrow |
| | | LB | Sensitive + regrow |
| | Liquid | AF | Sensitive + regrow |
| | | LB | Sensitive + regrow |
| ZLWE | Solid | AF | Resistant |
| | | LB | Resistant |
| | Liquid | AF | Resistant |
| | | LB | Resistant |
| WY3Z | Solid | AF | Sensitive |
| | | LB | Sensitive |
| | Liquid | AF | Sensitive |
| | | LB | Sensitive |

**Supplementary Table 1. Kinetic assay results are not affected by phage propagation method or media peptone source.** (LB=Luria Bertani, AF=animal-free)

| Phage | Isolate | Media Type | Kinetic Form |
|---|---|---|---|
| ACH8 | Original Host | AF | Sensitive + regrow |
| | | LB | Sensitive + regrow |
| | M1C086 | AF | Sensitive + regrow |
| | | LB | Sensitive + regrow |
| | M1C108 | AF | Sensitive + regrow |
| | | LB | Sensitive + regrow |
| WY3Z | Original Host | AF | Sensitive |
| | | LB | Sensitive |
| | SA01 | AF | Resistant |
| | | LB | Resistant |

**Supplementary Table 2. Peptone source of phage propagation media does not alter phage screen kinetic results.** (LB=Luria Bertani, AF=animal-free)

| Isolate | Media type | Number of reads | Total bases (Mb) | Median length (bp) | Read N50 | Median quality |
|---|---|---|---|---|---|---|
| M1C014 | AF | 833544 | 6044 | 5270 | 9284 | 21.3 |
| | LB | 236574 | 2500 | 6841 | 16588 | 21.5 |
| M1C034 | AF | 375696 | 3556 | 6510 | 13828 | 21.4 |
| | LB | 226463 | 2070 | 6482 | 13060 | 21.3 |
| M1C037 | AF | 596912 | 3177 | 4105 | 6636 | 21.0 |
| | LB | 353198 | 3580 | 7538 | 14607 | 21.7 |
| M1C054 | AF | 449998 | 3352 | 5564 | 9719 | 21.0 |
| | LB | 385703 | 3043 | 6061 | 10448 | 21.2 |
| M1C066 | AF | 903398 | 6247 | 5606 | 8528 | 21.4 |
| | LB | 218228 | 2415 | 7752 | 16718 | 21.8 |
| M1C077 | AF | 561229 | 3506 | 4394 | 8235 | 21.1 |
| | LB | 425820 | 3842 | 6009 | 13497 | 22.7 |
| M1C090 | AF | 808808 | 5325 | 4406 | 9707 | 20.9 |
| | LB | 317740 | 2823 | 6189 | 13280 | 21.2 |
| M1C093 | AF | 841420 | 5470 | 5162 | 8086 | 21.2 |
| | LB | 333956 | 2975 | 6207 | 12410 | 21.4 |
| M1C141 | AF | 411061 | 2114 | 3501 | 7304 | 20.6 |
| | LB | 373475 | 3296 | 6042 | 12979 | 21.4 |
| PAO1 | AF | 198334 | 1648 | 5987 | 11338 | 21.6 |
| | LB | 185048 | 1916 | 6861 | 16145 | 21.8 |
| SA01 | AF | 197008 | 2462 | 8953 | 18365 | 22.1 |
| | LB | 188333 | 2254 | 8139 | 18207 | 22.0 |
| SA09 | AF | 195145 | 2469 | 8916 | 19133 | 22.2 |
| | LB | 168344 | 1990 | 8377 | 17460 | 22.1 |

**Supplementary Table 3. Bacterial whole genome sequencing read metrics.** (LB=Luria Bertani, AF=animal-free)

| Phage | Media type | Number of reads | Total bases (Mb) | Median length (bp) | Read N50 | Median quality | Library Preparation |
|---|---|---|---|---|---|---|---|
| ACH8 | AF | 16073 | 178.4 | 7610 | 15461 | 18.2 | Native Barcoding |
| | LB | 30779 | 302.6 | 8015 | 12834 | 18.3 | |
| CEIU | AF | 8284 | 91.9 | 6971.5 | 18247 | 17.1 | Native Barcoding |
| | LB | 5281 | 74.6 | 7591 | 29929 | 17.1 | |
| FJBP | AF | 821727 | 2898.7 | 3524 | 3855 | 18.5 | Rapid PCR Barcoding |
| | LB | 610010 | 2384.2 | 3888 | 4314 | 18.7 | |
| K100 | AF | 1222708 | 4604.4 | 3728 | 4150 | 18.7 | Rapid PCR Barcoding |
| | LB | 629223 | 2767.6 | 4369 | 4884 | 18.9 | |
| KMBS | AF | 24192 | 123.0 | 3505.5 | 7139 | 17.9 | Native Barcoding |
| | LB | 55009 | 323.2 | 4108 | 8352 | 18.3 | |
| N0M1 | AF | 1697474 | 6904.0 | 4077 | 4435 | 18.9 | Rapid PCR Barcoding |
| | LB | 454699 | 1741.4 | 3797 | 4230 | 18.7 | |
| NYK5 | AF | 55009 | 323.2 | 4108 | 8352 | 18.3 | Native Barcoding |
| | LB | 10851 | 115.7 | 8785 | 14195 | 17.9 | |
| WY3Z | AF | 31322 | 268.0 | 6876.5 | 11634 | 18 | Native Barcoding |
| | LB | 49725 | 446.4 | 6893 | 12502 | 17.8 | |
| ZLWE | AF | 6844 | 163.3 | 19164.5 | 44237 | 18.5 | Native Barcoding |
| | LB | 19102 | 426.4 | 16488.5 | 44242 | 18.6 | |

**Supplementary Table 4. Phage whole genome sequencing read metrics.** (LB=Luria Bertani, AF=animal-free)

| Phage | Position | Ref (LB) | Alt (AF) | Total depth | Depth (Ref) | Depth (Alt) | Frequency | Location |
|---|---|---|---|---|---|---|---|---|
| FJBP | 51375 | C | T | 1847 | 338 | 1499 | 0.8116 | hypothetical protein |
| K100 | 51382 | C | T | 1648 | 288 | 1358 | 0.824 | hypothetical protein |
| NYK5 | 68455 | TG | T | 1266 | 27 | 1217 | 0.9613 | hypothetical protein |
| WY3Z | 38147 | A | T | 317 | 43 | 271 | 0.8549 | virion structural protein |
| | 72264 | AT | A | 347 | 156 | 186 | 0.536 | hypothetical protein |

**Supplementary Table 7. Phage SNPs.** A=adenine, C=cytosine, T=thymine, G=guanine
